# LRP2 controls sonic hedgehog-dependent differentiation of cardiac progenitor cells during outflow tract formation

**DOI:** 10.1101/801910

**Authors:** Annabel Christ, Thomas E. Willnow

## Abstract

Conotruncal malformations are a major cause of congenital heart defects in newborn infants. Recently, genetic screens in humans and mouse models have identified mutations in *LRP2* as a novel cause of a common arterial trunk, a severe form of outflow tract (OFT) defect. Yet, the underlying mechanism why the morphogen receptor LRP2 is essential for OFT development remained unexplained. Studying LRP2-deficient mouse models, we now show that LRP2 is expressed in the cardiac progenitor niche of the anterior second heart field (SHF) that contributes to elongation of the OFT during separation into aorta and pulmonary trunk. Loss of LRP2 in mutant mice results in depletion of a pool of sonic hedgehog-dependent progenitor cells in the SHF due to premature differentiation into cardiomyocytes as they migrate into the OFT myocardium. Depletion of this cardiac progenitor cell pool results in aberrant shortening of the OFT, the cause of CAT formation in affected mice. Our findings identified the molecular mechanism whereby LRP2 controls maintenance of progenitor cell fate in the anterior SHF essential for OFT separation, and why receptor dysfunction is a novel cause of conotruncal malformation.

## INTRODUCTION

LRP2 is a member of the LDL receptor gene family, a class of multifunctional endocytic receptors that play key roles in embryonic development and that cause severe developmental malformations in humans and animal models when dysfunctional (1). During neurulation, LRP2 is prominently expressed in the neuroepithelium that gives rise to the various parts of the developing central nervous system. Loss of receptor expression in this tissue in gene-targeted mice results in fusion of the forebrain hemispheres (holoprosencephaly) (2,3) and in overgrowth of the eye globe (buphthalmia) (4,5). Similar defects are seen in patients with Donnai-Barrow/ Facio-oculo-acoustico-renal (DB/FOAR) syndrome, an autosomal recessive disorder caused by inheritable *LRP2* mutations (6–9).

Concerning its mode of action, LRP2 has been shown to act as an auxiliary receptor for sonic hedgehog (SHH) and to activate or inhibit this morphogen pathway dependent on the biological context. In the neuroepithelium, LRP2 acts as a co-receptor to Patched1 (PTCH1) to promote SHH signaling and to pattern the ventral midline of the forebrain (10). By contrast, in the developing eye, it operates as a clearance receptor for SHH to antagonize growth promoting signals by this morphogen in the retina (11). Conceptually, these molecular functions of LRP2 in control of SHH signaling explain the forebrain and eye phenotypes observed in patients with DB/FOAR syndrome. However, LRP2 has also been shown to bind fibroblast growth factor (FGF) 8 (12) and bone morphogenetic protein (BMP) 4 (3), and may thus have the potential to regulate multiple morphogen pathways during organogenesis.

Surprisingly, unbiased screens using exome sequencing now implicated mutations in *LRP2* in congenital heart disease in humans, a receptor function not considered thus far (13). Additional evidence for a role in heart development came with ENU-induced mutagenesis studies that revealed *Lrp2* mutations as a prominent cause of cardiac outflow tract (OFT) defects in mice (14). The cardiac OFT is a transient structure at the arterial pole of the embryonic heart. During development, it separates into the ascending aorta and pulmonary trunk, outlets of the definitive left and right ventricle respectively. Defects in OFT formation produce conotruncal malformations characterized by incomplete or absent septation of the aorta and pulmonary trunk, resulting in low oxygen supply due to provision of mixed blood to the circulation. OFT malformations account for almost 30% of all congenital heart defects in humans (15). Recently, conotruncal malformations were also documented in mice with targeted *Lrp2* gene disruption, but the mechanism of receptor action remained unexplained (16).

We now demonstrate that LRP2 is specifically expressed in SHH-responsive progenitor cells in the dorsal pericardial wall (DPW) that contribute to formation of the OFT. Loss of receptor activity in *Lrp2* mutant mice impairs SHH signaling in the DPW, resulting in decreased numbers and disturbed myocardial differentiation of progenitors that migrate into the OFT myocardium. Ultimately, these morphogenetic defects cause insufficient elongation of the OFT, the reason for the conotruncal malformations seen in mice, and possible patients lacking the morphogen receptor LRP2.

## RESULTS

### Loss of LRP2 results in formation of a common arterial trunk (CAT) due to defective endocardial cushion formation

The OFT, positioned above the right ventricle, connects the embryonic ventricles with the aortic sac. During heart tube elongation and heart looping, the single OFT vessel is remodeled to undergo septation. Septation results in formation of the ascending aorta (Ao) from the left ventricle and the pulmonary trunk (Pa) arising from the right ventricle and ensures delivery of oxygenated blood to the body but deoxygenated blood to the lungs. To clarify the impact of LRP2 deficiency on OFT formation, we studied mice homozygous for a targeted *Lrp2* gene disruption (*Lrp2*^*−/−*^) generated by us previously (2). LRP2 mutant embryos were compared to their respective wild-type or heterozygous littermates, as no defects were observed in *Lrp2*^*+/−*^ animals. The latter two genotypes are jointly referred to as controls herein. Loss of LRP2 activity in *Lrp2*^*−/−*^ mice lead to defects in OFT formation characterized by incomplete or absent septation of the Ao and Pa, a defect referred to as a common arterial trunk (CAT; persistent truncus arteriosus). CAT formation was evident at embryonic (E) day 15.5 and 18.5 when control embryos showed properly separated Ao and Pa (Fig. 1A), while *Lrp2*^*−/−*^ embryos exhibited a single vessel exiting the heart from the right ventricle (Fig. 1A, asterisks). The above phenotype was seen in mice on an inbred C57BL/6N background. Interestingly, when we investigated the phenotype caused by loss of LRP2 function in *Lrp2*^*+/−*^ animals crossed with the ENU line 267 that carries a premature stop codon in the sequence encoding the extracellular domain of LRP2 (*Lrp2^+/267^*), compound heterozygous mutant mice (*Lrp2*^−/267^, on a mixed C57Bl/6N/FVBN genetic background) either developed a milder form of OFT defects, termed a double outlet right ventricle (DORV; 17 out of 33; Fig. S1), or showed normal OFT development with nicely separated Ao and Pa (16 out of 33; Fig. S1). This observation suggested genetic modifiers to impact LRP2 function in heart morphogenesis.

**Figure 1.**
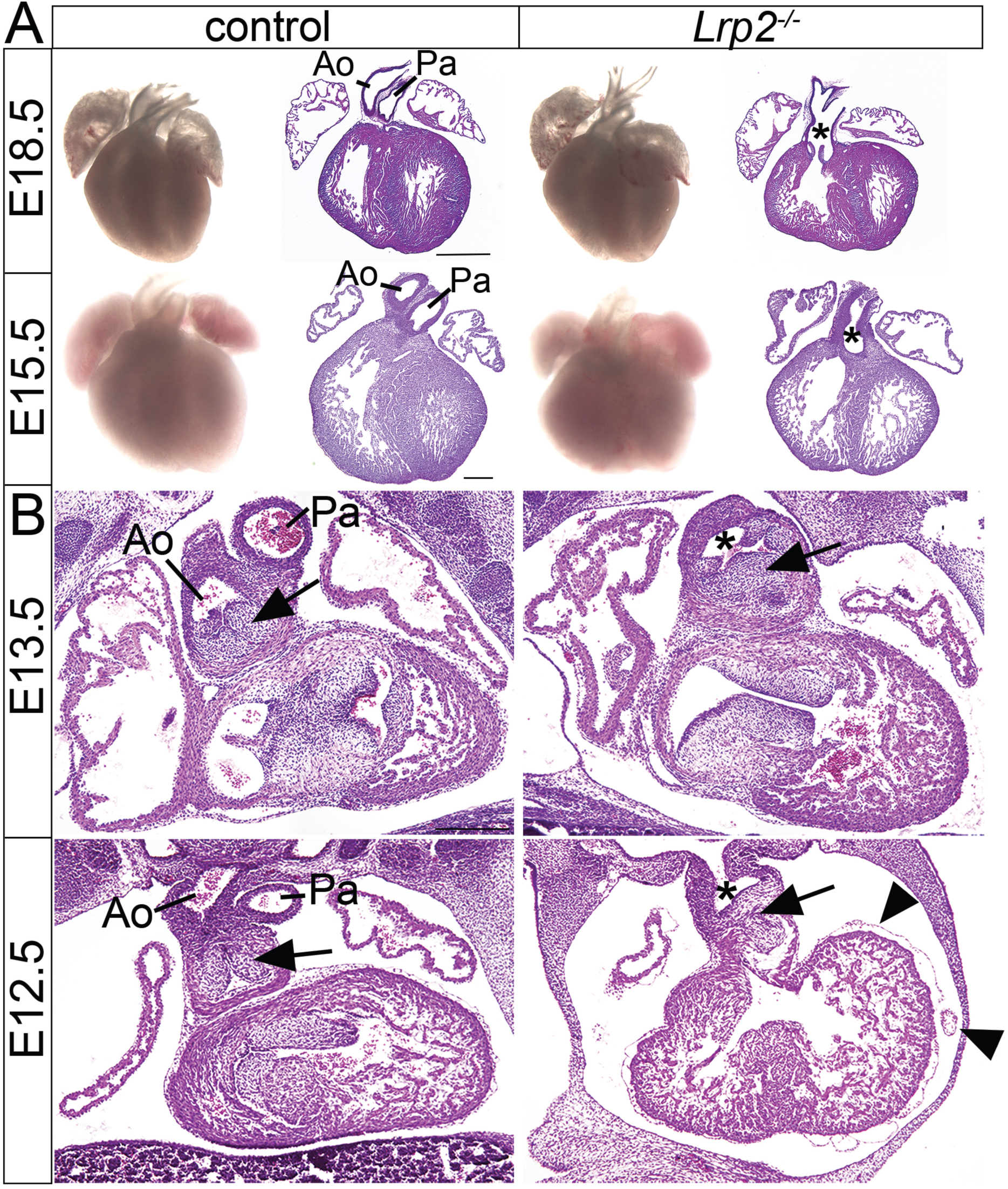
Loss of LRP2 causes formation of a common arterial trunk. (**A**) Whole mounts (left) and coronal sections (right) of *Lrp2*^*−/−*^ and control mouse hearts at embryonic (E) day 18.5 and 15.5. In control embryos, separation of the outflow tract into aorta (Ao) and pulmonary trunk (Pa) is evident in both embryonic stages. In *Lrp2*^*−/−*^ embryos, aorta and pulmonary trunk fail to separate resulting in a common arterial trunk (CAT, indicated by asterisks). This CAT phenotype showed full penetrance and was seen in 23 out of 23 *Lrp2*^*−/−*^ embryos analyzed. (**B**) Coronal sections of control and *Lrp2*^*−/−*^ hearts at E13.5 and E12.5. In control embryos, aorta (Ao) and pulmonary trunk (Pa) are present. Arrows indicate two distinct swellings forming the endocardial cushions in the aortic valve in control hearts. In *Lrp2*^*−/−*^ hearts, a common outflow vessel (indicated by asterisks) above the right ventricle is formed. Instead of two endocardial swellings only one cell cluster in endocardial cushion formation (arrow) is apparent. Arrowheads indicate blebbing of the epicardium and hemorrhages in *Lrp2*^*−/−*^ embryos at E12.5. Scale bars: A, 750 µm; B, 250 μm

In the following, we focused our studies on *Lrp2* mutants on a pure C57BL/6N genetic background exhibiting CAT formation. Histological alterations indicative of an OFT defect manifested around E12.5 to E13.5 when OFT septation normally occurs. Separation of Ao and Pa through the formation of endocardial cushions was seen in control embryos (Fig 1B). By contrast, LRP2-deficient embryos failed to form distinct endocardial cushions but exhibited an unorganized cell cluster in the OFT (Fig. 1B, arrows). In addition, blebbing of the epicardial layer as well as hemorrhages were visible in mutant hearts (Fig. 1B, arrowheads).

We further investigated defects in endocardial cushion formation in *Lrp2*^*−/−*^ embryos by *in situ* hybridization (ISH) for *Sox9* (Fig. 2A). Sox9 is expressed in the cardiac cushion mesenchyme and promotes cardiac precursor cell expansion during heart valve development. At E10.5, two *Sox9* positive endocardial cushions were visible in control embryos (arrows) while these distinct tissue swellings were not detectable in *Lrp2*^*−/−*^ embryos, despite the presence of *Sox9* positive cells. By E11.5, these spiraling clusters of *Sox9* positive cells (arrows) were less compacted in *Lrp2*^*−/−*^ compared to control embryos and failed to develop into the endocardial swellings of the aortic valve (E12.5, stippled circles in controls). Rather, they formed an unorganized *Sox9* positive cell cluster (E12.5, stippled circle in *Lrp2*^*−/−*^). Failure of the development of the endocardial cushions lead to a malformed outlet septum (Fig. 2B, asterisk) that subsequently resulted in the formation of a CAT in LRP2-deficient embryos.

**Figure 2.**
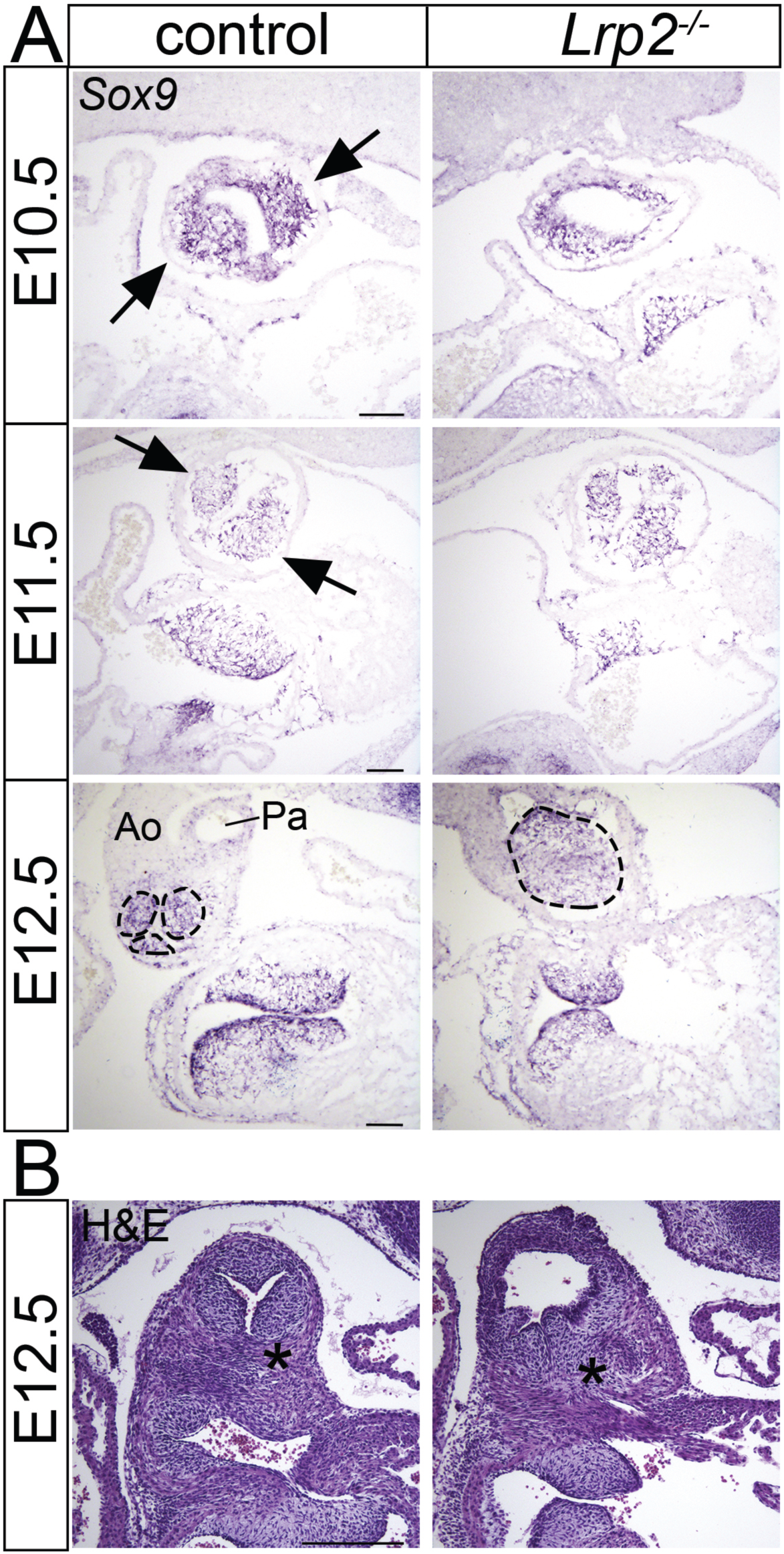
Altered endocardial cushion formation in *Lrp2*^*−/−*^ hearts. **(A)** *In situ* hybridization (ISH) for *Sox9* on coronal sections of *Lrp2*^*−/−*^ and control embryos depict endocardial cushion formation in the OFT. At E10.5, two endocardial cushions are formed by two distinct tissue swellings in the control OFT tissue (arrows). In *Lrp2*^*−/−*^ OFT tissue, these two tissue swellings are not visible. At E11.5, *Sox9* positive cells cluster in the endocardial cushions (arrows) in control OFT tissue. In mutants, *Sox9* positive cells are less compacted and poorly organized in the endocardial cushions. At E12.5 in control OFT tissue, the OFT septum has formed and divides aorta (Ao) and pulmonary trunk (Pa). *Sox9* positive cells are localized in the endocardial swellings of the aortic valve (stippled circles), while in *Lrp2*^*−/−*^ OFT vessel only a disorganized cluster of *Sox9* positive cells is visible (stippled circle). **(B)** Hematoxilin and eosin-stained coronal sections of *Lrp2*^*−/−*^ and control embryos documenting endocardial cushions in the aortic valve of control and *Lrp2*^*−/−*^ embryos. While the aorticopulmonary septum has formed in control embryos (asterisk), it failed to properly develop in *Lrp2*^*−/−*^ embryos (asterisk). Scale bar: 100 µm

### LRP2 is expressed in the progenitor domain of the second heart field

The formation, elongation, and septation of the cardiac OFT depends on the interaction of two distinct cell populations, namely cardiac neural crest cells (CNCC) and second heart field (SHF) cells. CNCC are a subpopulation of neural crest cells that originate from the dorsal neural tube and migrate into pharyngeal arches 3, 4, and 6. Starting from E9.5, CNCC populate the cardiac outflow tract and migrate into the outflow tract cushions (17). There, they give rise to the condensed mesenchyme in the endocardial cushions forming the aorticopulmonary septation complex that divides the distal OFT into Ao and Pa (18). SHF cells are cardiac progenitor cells located in the pharyngeal mesoderm. During heart development they are added to the arterial pole of the OFT driving OFT elongation.

To interrogate the role of LRP2 during OFT formation, we investigated the expression pattern of the receptor during OFT separation using immunohistology. At E10.5, when CNCCs invaded the OFT cushions, LRP2 showed strong expression in the surface ectoderm as well as in the epithelium of the pericardial cavity, including the dorsal pericardial wall (DPW) harboring SHF progenitor cells with atypical apicobasal polarization. In addition, LRP2 expression was detected in the epithelium lining the distal outflow tract (OFT) (Fig. 3A, E10.5). Upon closer inspection of the distal OFT (Fig. 3B, coronal view), LRP2 localized mainly to the superior and inferior regions of the OFT. LRP2 immunoreactivity was enriched in the OFT wall overlying the aortic (AoIC) and pulmonary intercalated cushions (PuIC). To better visualize LRP2 expression in the DPW, we used higher magnification of a sagittal view of E10.5 control hearts (Fig. 3B, sagittal). This plane of section demonstrated that LRP2 is abundantly expressed in the DPW while levels are reduced in the transition zone (Tz) and completely absent from the OFT myocardium. At E12.5, when the OFT in controls was already septated, robust LRP2 expression persisted in the DPW and in myocardial cells in the most distal regions of the Ao and Pa, while only minor expression levels were visible in atria and ventricles (Fig. 3A; E12.5; arrowheads). Based on these data, we concluded that LRP2 mainly localizes to the SHF with highest expression levels in the SHF progenitor zone of the DPW.

**Figure 3.**
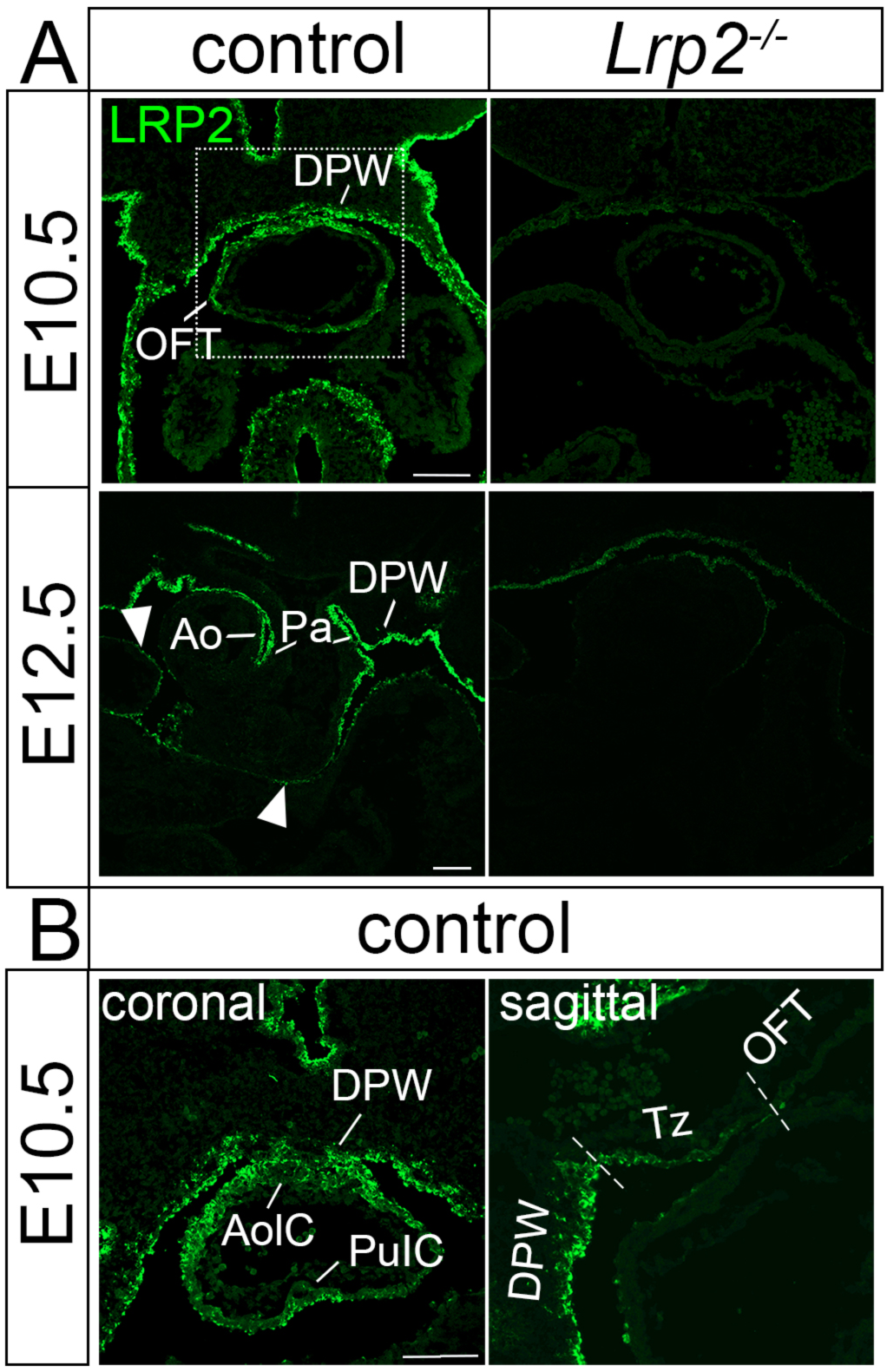
LRP2 expression in the developing outflow tract. Immunhistological detection of LRP2 on coronal sections of the embryonic mouse heart at the indicated embryonic days. (**A**) At E10.5, LRP2 expression is seen along the ectoderm as well as in the epithelium enclosing the pericardial cavity, including the dorsal pericardial wall (DPW) and in the distal outflow tract (OFT) myocardium of controls. No immunoreactivity for LRP2 is detected in receptor mutant hearts. At E12.5, LRP2 expression is visible in the DPW and in myocardial cells lining the pulmonary artery (Pa) as well as the side of the aorta (Ao) facing the Pa. Minor LRP2 levels are visible in the epithelium lining atria and ventricles (arrowheads). (**B**) Higher magnification of the region corresponding to the boxed area in the control embryos in panel A. On a coronal view (left panel), robust LRP2 expression is seen in cells of the second heart field in the dorsal pericardial wall (DPW) and in distal myocardial cells of the outflow tract (OFT). In the OFT, LRP2 immunoreactivity is most prominent in the OFT wall overlying the swellings of the aortic (AoIC) and pulmonary intercalated cushions (PuIC). On a sagittal view (right panel), prominent levels of LRP2 are seen in epithelial cells of the DPW. Receptor expression is significantly reduced in the transition zone (Tz) and completely absent from the proximal OFT myocardium (OFT). Boundaries between DPW, TZ, and OFT myocardium are marked by stippled white lines. Scale bar: 100 μm

### LRP2 deficiency specifically impacts second heart field progenitor cells

While LRP2 expression in neural crest cells had been reported before (19, 20), we failed to detect the receptor in the CNCC population by immunohistology (Fig. 4A). Still, the endocardial cushion defect observed in *Lrp2*^*−/−*^ embryos (Fig. 2) was consistent with a potential defect in the CNCC population. To query an indirect effect of receptor deficiency on this cell population, we crossed the *Lrp2* deficient mouse strain with the transgenic *Wnt1-Cre_LacZ* reporter line to specifically mark CNCCs (21). Staining for lacZ activity at E10.5 revealed a comparable pattern of CNCCs in the OFT in *Lrp2*^*−/−*^ and control embryos (Fig. 4B). Subtle differences were seen in more proximal regions of the spiraling OFT cushions, where CNCCs appeared less compacted in *Lrp2*^*−/−*^ embryos compared with controls (Fig. 4B, stippled circle). We concluded that LRP2 deficiency did not affect the migration of CNCCs into the OFT and their integration in the endocardial cushions. Rather subtle alterations in CNCC organization seen in the proximal OFT of mutant mice were considered secondary to defects in cushion formation described in Fig. 2.

**Figure 4.**
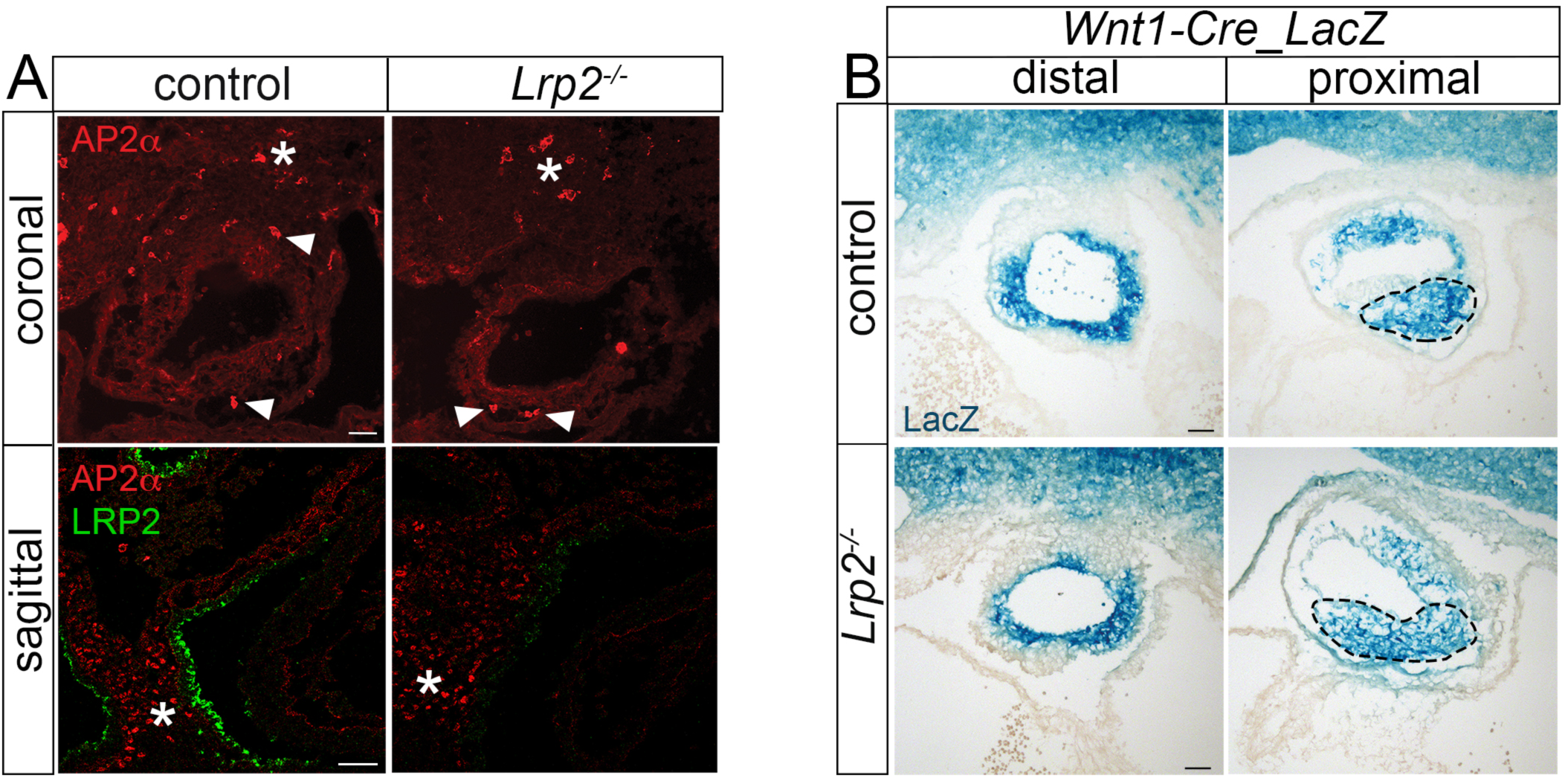
LRP2 deficiency does not affect the cardiac neural crest cell (CNCC) population migrating into the OFT. (**A**) Immunohistological detection of AP2α on coronal and of AP2α and LRP2 on sagittal sections of the hearts of control and *Lrp2*^*−/−*^ embryos at E10.5. Similar numbers of AP2α positive cells are detected in the second heart field region (asterisks) and in the OFT (arrowheads) of *Lrp2*^*−/−*^ and control embryos. AP2α and LRP2 show non-overlapping expression domains, suggesting absence of LRP2 from CNCCs. (**B**) Staining for lacZ activity in CNCCs on coronal sections of E10.5 mouse OFT tissues from the *Wnt1-Cre_LacZ* reporter line. Mice express (control) or lack LRP2 (*Lrp2*^*−/−*^). In sections of the distal OFT region (left panels), the presence of CNCCs in the outflow vessel is detected in control and *Lrp2*^*−/−*^ embryos. In more proximal regions of the OFT (right panel), CNCCs fail to accumulate in the endocardial cushions in *Lrp2*^*−/−*^ as compared with the control OFT vessel (stippled circle). Scale bars: 50 µm

Since *Lrp2* was not expressed in CNCC and LRP2 deficiency did not affect the migration of CNCC into the heart, we turned our attention to SHF cells that express LRP2 and are necessary for OFT septation. Among other markers, these cells are characterized by expression of insulin gene enhancer protein (*Islet1*) (22). Detection of Islet1 protein as well as *Islet1* transcript (Fig. 5A and B) showed decreased expression levels in the pharyngeal mesoderm and in the OFT in *Lrp2*^*−/−*^ embryos compared with controls. In the OFT, Islet1 protein and transcript were especially reduced in the superior and inferior intercalated cushions (Fig. 5A and B, arrowheads). Reduced levels of Islet1 in the intercalated cushions may explain the defect in endocardial cushion formation seen in the aortic valve of *Lrp2*^*−/−*^ embryos (Fig. 2).

**Figure 5.**
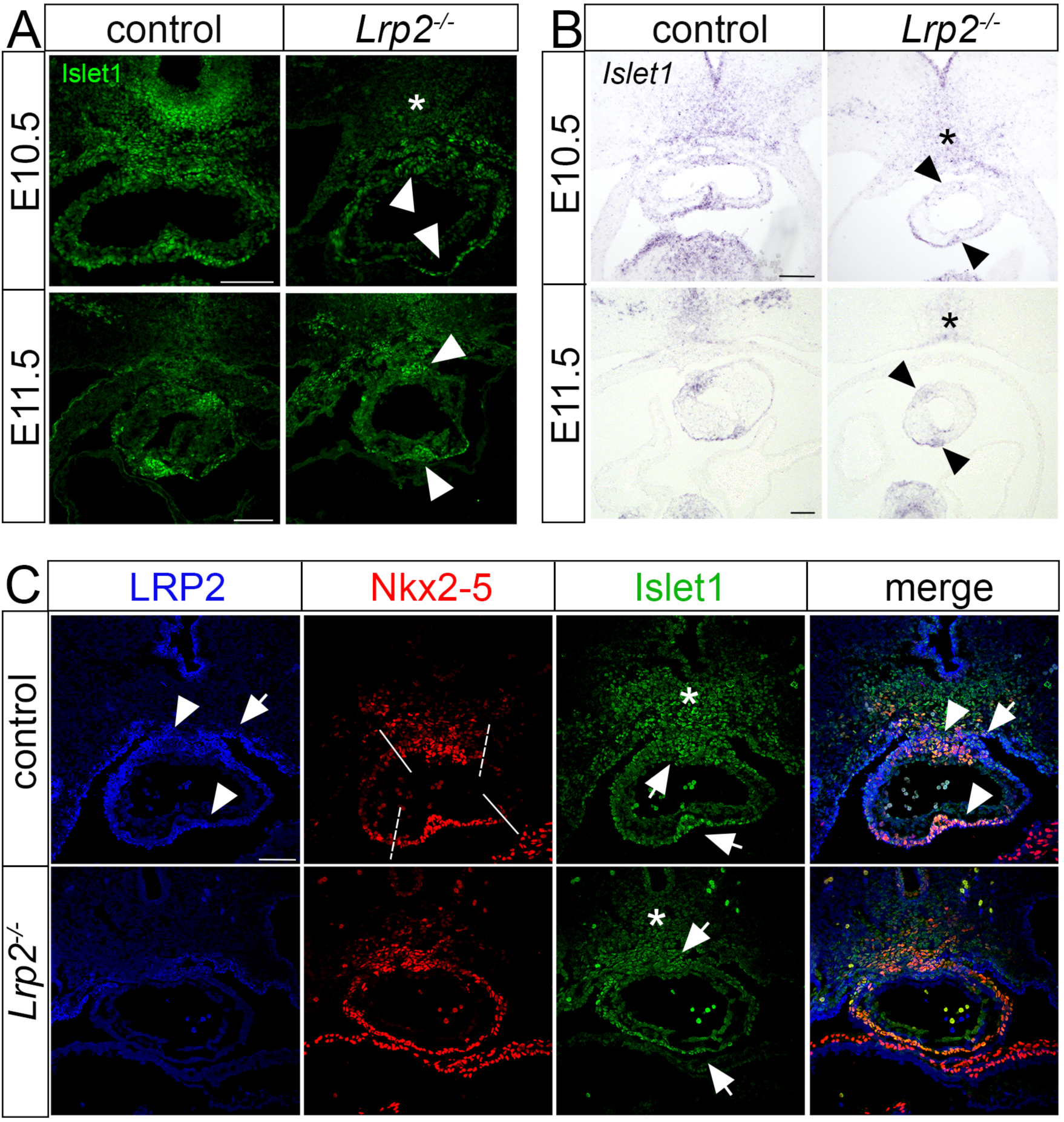
LRP2 deficiency causes a reduction in second heart field (SHF) cardiac progenitor cells. (**A**) Immunodetection of Islet1 and (**B**) *in situ* hybridization for *Islet1* on coronal OFT sections of control and *Lrp2*^*−/−*^ embryos at E10.5 and E11.5. In *Lrp2*^*−/−*^ embryos, immunosignals for Islet1 are decreased in the pharyngeal mesoderm (A, asterisk) and in the superior and inferior intercalated cushions of the OFT (A, arrowheads) when compared to controls. The reduction in Islet1 protein levels in mutants is paralleled by decreased *Islet1* transcript levels at both embryonic stages in the pharyngeal mesoderm (B, asterisk) and in the intercalated cushions (B, arrowheads) as compared with controls. Scale bar: 100 µm. (**C)** Immunohistological detection of LRP2, of second heart field (SHF) marker Nkx2-5 and of SHF cardiac progenitor marker Islet1 on coronal sections of E10.5 control and *Lrp2*^*−/−*^ embryos. Single channels as well as merged images are shown. Cardiac progenitor cells of the SHF in the dorsal pericardial wall (arrow in LRP2 and merged channels) and in the myocardium lining the intercalated cushions (arrowheads in LRP2 and merged channels) stain positive for LRP2. Lack of LRP2 in *Lrp2*^*−/−*^ embryos results in the presence of Nkx2-5 positive cells throughout the OFT. This contrasts with the more restricted pattern in the superior and inferior OFT region (stippled white lines) in the control embryo. Furthermore, *Lrp2*^*−/−*^ embryos show reduced numbers of Islet1 positive cells in the pharyngeal mesoderm (asterisk in single channel) as well as in the intercalated cushions (arrows in single channel) when compared with controls. Scale bar: 75 µm

To investigate in more detail which cells within the SHF express LRP2 we investigated the localization of LRP2 with respect to SHF marker Islet1 and cardiac mesoderm marker NK2 transcription factor related, locus 5 (Nkx2-5), a myocardial transcription factor regulated by Islet1 and required for SHF development. LRP2 was expressed in the DPW (Fig. 5C, arrow) and in the myocardium lining the intercalated cushions in the OFT (Fig. 5C, arrowheads; and Fig. 3B), partially overlapping in these regions with Nkx2-5 and Islet1 in controls. In *Lrp2*^*−/−*^ mutants, Nkx2-5 protein in the distal OFT was less restricted to the superior and inferior regions (Fig. 5C, stippled lines in control embryos) but showed a more widespread localization throughout the distal OFT. Islet1 demonstrated decreased levels in the pharyngeal mesoderm (Fig. 5A and C, asterisks) as well as in the superior and inferior intercalated cushions (Fig. 5A arrowheads; Fig. 5C arrows). We also tested expression of two additional SHF markers, namely neurovascular guiding factor *Sema3c*, important to correctly navigate CNCCs towards the OFT (23), as well as *T-box transcription factor* (*Tbx*) (Fig. S2). Despite an overall unchanged expression pattern of *Sema3c*, slightly reduced levels in the intercalated cushions of the OFT were seen on coronal sections of *Lrp2*^*−/−*^ embryos compared with controls (Fig. S2A, arrowheads). By contrast, expression levels of *Tbx1* (Fig. S2B) were comparable between genotype groups suggesting mis-patterning of a specific population of SHF progenitors rather than complete absence of SHF progenitors in LRP2 mutant mice.

### LRP2 promotes SHH signaling during outflow tract formation

So far, our data documented a decrease in Islet1 positive SHF progenitor cells and a shifted localization of Nkx2-5 positive cells in LRP2 mutant embryos, arguing for a role of this receptor in promoting cardiac progenitor maintenance during OFT septation. Several signaling pathways have been implicated in the formation, proliferation, and differentiation of multipotent cardiac progenitor cells in the SHF (24). Since LRP2 is known to act in multiple morphogen pathways, we investigated these pathways in *Lrp2*^*−/−*^ embryos during OFT development.

Canonical Wnt signaling has been shown to act upstream of other signaling pathways in control of Islet1 positive progenitor cell proliferation (25). Non-canonical Wnt signaling is implicated in OFT development (26, 27) and important for planar cell polarity during myocardialization of the OFT cushions. It regulates polarity and intracellular cytoskeletal rearrangements, important to prevent premature differentiation of SHF progenitor cells into cardiomyocytes (28–30). To explore potential defects in canonical Wnt signaling during OFT formation, we crossed the LRP2 mutant strain with the *Tcf/Lef_LacZ* reporter line. Detecting lacZ activity as a read-out of Wnt signaling, we failed to observe any changes in Wnt activity in the SHF of embryos lacking LRP2 (Fig. S3A). Also, the non-canonical Wnt pathway was unchanged as deduced from investigating the expression pattern of *Wnt11* in the OFT using ISH (Fig S3B). BMP signaling downregulates proliferation when SHF cells enter the OFT and it drives myocardial differentiation (31). Analyzing the BMP pathway by studying *Bmp4* expression in the distal OFT failed to detect obvious changes in LRP2-deficient as compared with control embryos (Fig. S3B).

Another essential morphogenetic signal in SHF patterning is provided by SHH secreted by the pharyngeal endoderm. SHH regulates both OFT development and CNCC survival. In addition, in the pharyngeal endoderm, SHH signaling coordinates a secondary, so far unknown pathway to control SHF survival and development (32). Using the *Gli1-LacZ* reporter allele introduced into the *Lrp2* mutant strain, we detected reduced SHH activity in the distal OFT of *Lrp2*^*−/−*^ embryos as compared with controls. This defect was especially evident in the inferior OFT region where SHH signaling was almost absent in LRP2-deficient embryos (Fig. 6, asterisks). To test if reduced SHH signaling in *Lrp2*^*−/−*^ embryos was connected to the SHF defects characterized by reduced *Islet1* expression, we crossed the LRP2-deficient line with the *Gli1-CreER^T2^* reporter strain. In this strain, expression of a tamoxifen-inducible Cre transgene under control of the *Gli1* promoter enables labeling of SHH-responsive cells by Cre-induced expression of YFP. To induce YFP expression, pregnant mice were injected with tamoxifen at E7.5 and embryos were analyzed at E10.5. Immunodetection of YFP demonstrated SHH-responsive cells accumulating in the inferior OFT region in control embryos (Fig. 7A). In *Lrp2*^*−/−*^ embryos, YFP-positive cells were reduced about 50% in numbers and failed to accumulate in the inferior OFT region (Fig. 7A, arrowheads; Fig. 7B). Next, we performed co-immunostaining for YFP and Islet1 to test if SHF progenitor cells were responsive to SHH signals. In line with this assumption, YFP positive cells in the inferior OFT of control embryos co-immunostained with Islet1. *Lrp2*^*−/−*^ embryos showed a significant reduction in Islet1immunoreactvity in the pharyngeal mesoderm (Fig. 7A, asterisk) as well as in the superior and inferior OFT (Fig. 7A, arrowheads). Quantification of cardiac progenitor cells double positive for Islet1 and YFP substantiated a significant reduction of SHH-responsive Islet1 positive progenitor cells in the distal OFT of LRP2-deficient embryos (Fig. 7C).

**Figure 6.**
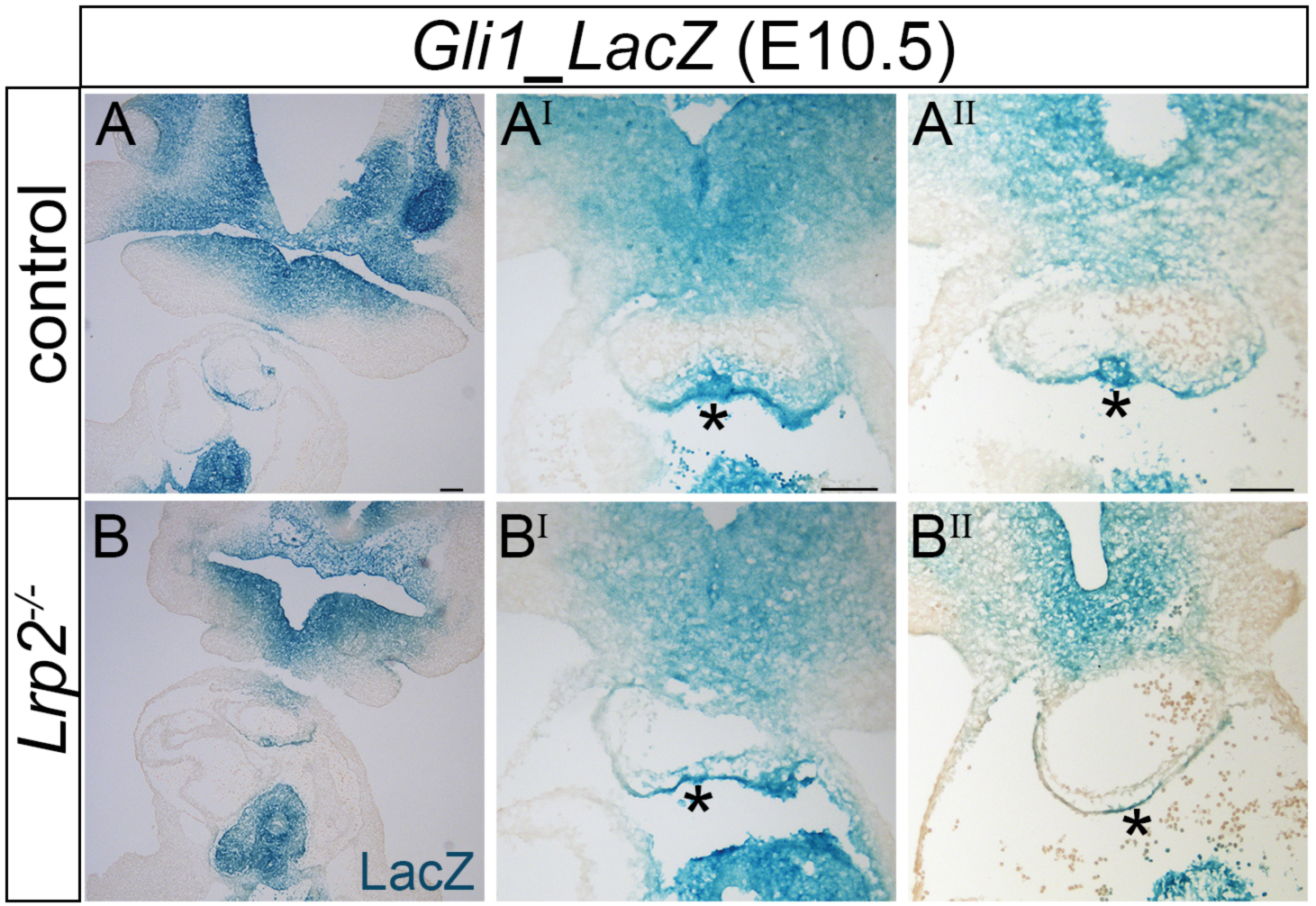
Reduced SHH signaling activity in the *Lrp2*^*−/−*^ OFT. Staining for lacZ activity on coronal sections from E10.5 *Gli1_LacZ* reporter mice to visualize SHH signaling. Animals express (control) or lack LRP2 (*Lrp2*^*−/−*^). Overview images of heart and pharyngeal region are shown in panels A and B. In addition, higher magnification images of two control (A^I^ - A^II^) and two mutant (B^I^ - B^II^) embryos are given. In *Lrp2*^*−/−*^ OFT vessel, the activity of the GLI1-dependent LacZ reporter is significantly reduced (compare asterisks in B^I^ to A^I^) or almost completely absent (compare asterisks in B^II^ to A^II^). Scale bar: 100 µm.

**Figure 7.**
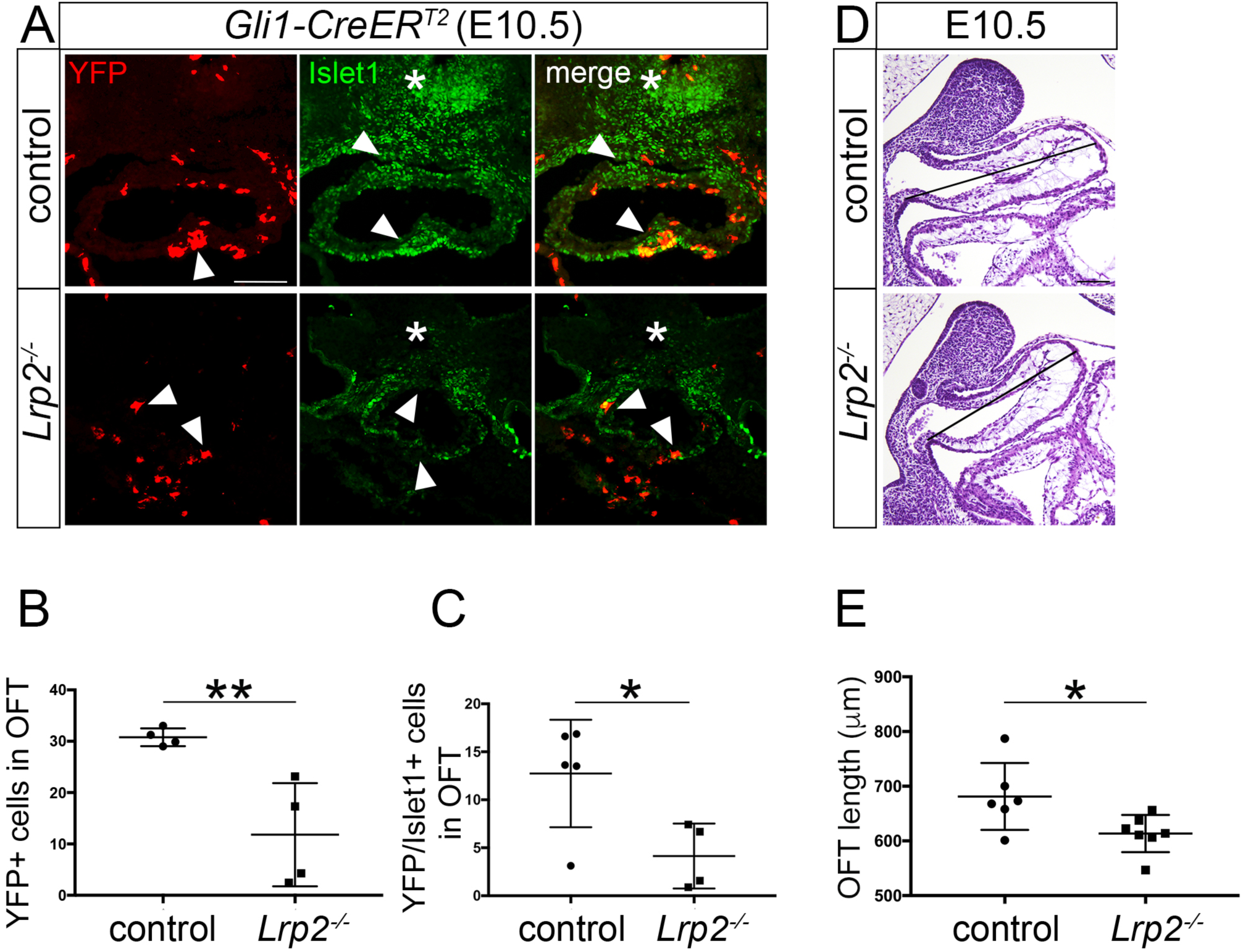
Reduced SHH responsiveness of cardiac progenitor cells in the *Lrp2*^*−/−*^ OFT. (**A**) Immunhistological detection of YFP and Islet1 on coronal sections of E10.5 mouse hearts from the tamoxifen-inducible *Gli1-CreER*^*T2*^ reporter strain. The animals express (control) or lack LRP2 (*Lrp2*^*−/−*^). Single channels as well as merged images are shown. In control embryos, YFP-labeled cells are visible throughout the OFT but accumulate in the inferior intercalated cushion (arrowhead). Islet1-positive cardiac progenitor cells are seen in the pharyngeal mesoderm (asterisk) as well as in the superior and inferior OFT (arrowheads). In *Lrp2*^*−/−*^ embryos, YFP positive cells fail to accumulate in the inferior OFT but are dispersed throughout the OFT vessel (arrowheads). Also, Islet1 positive cells are reduced in the pharyngeal mesoderm (asterisk) and in the OFT vessel (arrowheads). Scale bar: 100 µm. (**B**) YFP positive cells (as exemplified in panel A) were quantified on 5 - 9 coronal sections through the distal OFT vessel of 4 control and 4 *Lrp2*^*−/−*^ embryos. YFP positive cells are reduced about 50% in *Lrp2*^*−/−*^ embryos. (**C**) Cardiac progenitor cells double positive for Islet1 and YFP (as exemplified in panel A) were quantified on 5 - 9 coronal sections through the distal OFT vessel of 5 control and 4 *Lrp2*^*−/−*^ embryos. The numbers of YFP/Islet1 double positive cells in the OTF are significantly reduced in mutant as compared with control embryos (**, p≤0.01 *, p≤0.05; unpaired Student’s t test). (**D**) Exemplary H&E stained sagittal sections of *Lrp2*^*−/−*^ and control embryos at E10.5. The lines indicate OFT length. Scale bar: 100 µm. (**E**) OFT length in control and *Lrp2*^*−/−*^ OFT tissues as quantified on 5 sagittal sections each through the OFT vessel of 6 control and 7 *Lrp2*^*−/−*^ embryos. OTF length is significantly reduced in mutant as compared with control embryos (*, p≤0.05; unpaired Student’s t test).

During OFT formation, SHF progenitor cells located in the DPW move along a path of differentiation to the transition zone (Tz), where they initiate a myocardial gene expression program. They finally reach the OFT myocardium to fully differentiate into cardiomyocytes. By this movement, cardiac SHF progenitor cells contribute to the elongation of the OFT, necessary for correct alignment. Disturbances in this differentiation program result in a shortened OFT and can give rise to a CAT (33). A significant decrease in OFT length was seen in the hearts of *Lrp2*^*−/−*^ embryos compared with controls (Fig. 7D and E), arguing for a defect in progenitor movement in *Lrp2*^*−/−*^ embryos.

To investigate the consequences of reduced SHH signaling in DPW progenitors for their differentiation potential, we studied sagittal sections of *Gli1-CreER^T2^* control and LRP2-deficient embryos at E10.5. In controls, SHH-responsive cells (as deduced by YFP expression) concentrated in the DPW and distal Tz, a region overlapping with the LRP2 expression domain in the DPW (Fig. 8A). Almost no SHH-responsive cells were detected in the OFT myocardium of control embryos as evidenced by an absence of co-staining of YFP with the cardiomyocyte marker MF20 (Fig. 8A). By contrast, in *Lrp2*^*−/−*^ embryos, an increased number of SHH-responsive cells were co-stained by YFP and MF20 in the OFT myocardium (Fig. 8A, arrowheads). Quantification of YFP and MF20 double positive cells corroborated this finding and demonstrated a significant increase in YFP-MF20 double positive cells in the OFT myocardium in *Lrp2*^*−/−*^ embryos (Fig. 8B). Jointly, the reduction of YFP/Islet1 double positive cells in the distal OFT and the increased amount of YFP/MF20 double positive cells in the OFT myocardium in *Lrp2*^*−/−*^ embryos argued for a role of LRP2 in facilitating the response of SHF progenitor cells in the DPW to SHH. Such a response is required to control migratory movement and differentiation along their path to the OFT myocardium. Reduced responsiveness to SHH in the LRP2-deficient DPW was not due to impaired formation of primary cilia as documented by staining for the ciliary marker Arl13b in Islet1 positive progenitor cells in both genotype groups (Fig. S4).

**Figure 8.**
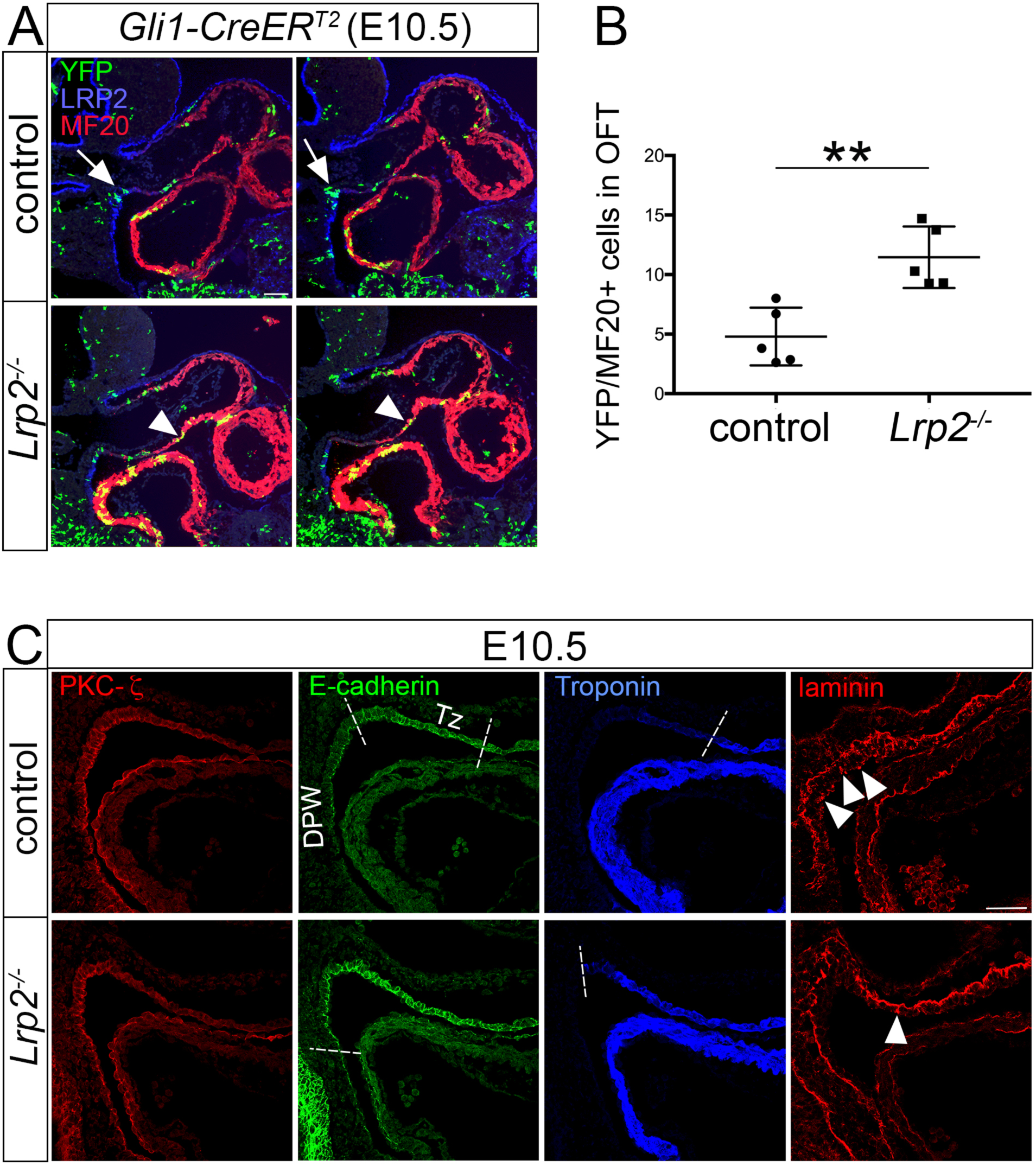
Loss of LRP2 decreases the SHF progenitor niche. **(A)** Immunodetection of YFP (green), LRP2 (blue), and MF20 (red) on two exemplary sagittal sections for E10.5 control and for *Lrp2*^*−/−*^ hearts of the *Gli1-CreER*^*T2*^ reporter line. In control embryos, SHH responsive (YFP positive) cells cluster at the border between dorsal pericardial wall (DPW) and transition zone (Tz; arrow). Very few double positive cells for YFP-MF20 are detected in the OFT of control embryos. In the two *Lrp2*^*−/−*^ heart tissue sections, MF20-positive cardiomyocytes in the OFT myocardium are positive for YFP (arrowhead), indicating aberrant differentiation of SHH responsive cells. Scale bar: 100 µm. **(B)** Cardiomyocytes double positive for MF20 and YFP (as exemplified in panel A) were quantified on 5 - 8 sagittal sections through the OFT vessel of 5 control and 5 *Lrp2*^*−/−*^ embryos. The numbers of YFP/MF20 double positive cells in the OTF are significantly reduced in mutant as compared with control embryos (**, p≤0.01; unpaired Student’s t test). **(C)** Immunohistological detection of PKC-ζ, E-cadherin, troponin, and laminin on sagittal sections of E10.5 control and *Lrp2*^*−/−*^ heart tissues. Signal intensity and distribution for adherens junction component PKC-ζ (red) are unchanged comparing control and *Lrp2*^*−/−*^ hearts. In control hearts, E-cadherin (green) exhibits strong expression in the transition zone (Tz) but reduced levels in the dorsal pericardial wall (DPW). In *Lrp2*^*−/−*^ hearts, the E-cadherin expression domain extends into the DPW (to the stippled line). Troponin (blue) marks differentiated cardiomyocytes in the OFT myocardium. In contrast to control hearts, troponin expression extends from the OFT myocardium into the transition zone (stippled line) in *Lrp2*^−/−^ hearts. Laminin, a marker for the basal lamina, shows a punctuated pattern with apical vesicles (arrowheads) in the DPW and distal Tz in controls. In *Lrp2*^*−/−*^ hearts, laminin-positive apical vesicles are reduced in numbers (arrowhead) and the regular pattern of the basal lamina from the OFT continues into the Tz and DPW. Scale bar: 50 µm

### LRP2 defines the boundary of the cardiac SHF progenitor zone

To further substantiate a role for LRP2 in control of SHF progenitor cell differentiation along a path from the DPW to the Tz and finally to the OFT myocardium, we performed immunodetection of markers representative for these three tissues in control and receptor mutant embryos. Previous studies demonstrated an atypical apicobasal polarization of SHF progenitor cells and their basal filopodia in the DPW, and their increasing epithelial characteristics in the Tz before differentiation in the OFT myocardium (29,34–36). Therefore, we focused on immunohistological detection of proteins having a role in apical polarity in *Lrp2*^*−/−*^ DPW and Tz compared with controls. Immunodetection of the apical tight junction marker aPKC-ζ (37) demonstrated robust expression at the apical cell membrane in the control DPW (Fig. 8C). No changes in signal intensity or localization in aPKC-ζ were detected in the *Lrp2*^*−/−*^ DPW, arguing against an overall effect of LRP2 deficiency on polarization of cells in the DPW. However, the situation was different for the adherens junction marker E-cadherin. Although localization of E-cadherin to the apical cell membrane was comparable between the two genotype groups, the distribution of E-cadherin-positive cells was clearly shifted in the mutants. As shown in Fig. 8C, in controls, E-cadherin was expressed in the anterior DPW and increased in expression in the Tz, consistent with an increasing epithelial cohesion before cells differentiate into myocardium in the OFT. Vanishing E-cadherin protein levels were detected in the posterior region of the DPW closer to the venous pole. By contrast, in *Lrp2*^*−/−*^ embryos, the localization of E-cadherin-positive cells in the DPW was shifted to more posterior regions in the DPW (Fig. 8C, compare localization of the stippled lines). This shifted cell identity was also confirmed by a shift in localization of cardiac troponin I, a marker for mature cardiomyocytes (Fig. 8C). Cardiac troponin I was exclusively expressed in cells of the OFT myocardium in controls. In *Lrp2*^*−/−*^ OFT tissue, also cells in the TZ stained positive for cardiac troponin I. Furthermore, laminin, a marker for the basal lamina showed an altered localization in *Lrp2*^*−/−*^ embryos. While in the control DPW and Tz, laminin showed a punctuated pattern in the apical cell region, this punctuated pattern was lost in the *Lrp2*^*−/−*^ DPW and Tz (Fig. 8C, arrowheads). In addition, in control embryos, the basal lamina formed a continuous pattern in the OFT region but not in the Tz, whereas in *Lrp2*^*−/−*^ hearts, this continuous pattern of the basal lamina reached inside the Tz (Fig. 8C). Collectively, these findings argue for a reduction in numbers and for premature differentiation of *Lrp2*^*−/−*^ SHF progenitor cells as they transit from the DWP via the Tz into OFT myocardium. These defects confound the contribution of SHF progenitor cells to OFT elongation and result in shortening of the OFT, the cause of CAT formation in *Lrp2*^*−/−*^ embryos.

## DISCUSSION

Previous studies implicated LRP2 in OFT formation (16). Still, the exact mechanism whereby this multifunctional endocytic receptor may control formation of the cardiac OFT and why receptor dysfunction results in conotruncal malformations in patients and in mouse models remained unexplained. Here, we uncovered a crucial role for LRP2 in SHH signaling in progenitor cells of the DPW, signals required to ensure their timely differentiation into mature cardiomyocytes as they migrate into the OFT tissue. Loss of LRP2 results in a decrease in SHH-responsive progenitor cells in the DPW and in a concomitant appearance of cardiomyocytes in the Tz, arguing for a depletion of this progenitor niche due to premature differentiation of cardiac progenitors along their path to an OFT myocardial fate.

### Loss of SHF progenitors is the cause of CAT formation in *Lrp2*^*−/−*^ embryos

Formation and subsequent septation of the OFT is dependent on two cell populations, the SHF and the CNCCs. We traced the primary defect in LRP2 deficiency to the progenitor cells of the SHF that express this receptor (Fig. 5C). While the CNCC population normally migrated into the OFT and reached the distal OFT cushions (Fig. 4), the number of Islet1 positive progenitor cells in the SHF was significantly reduced in receptor mutant hearts (Fig. 5). Cardiac progenitors in the DPW contribute to growth and elongation of the OFT, a process essential for proper septation into Ao and Pa (33). *Lrp2*^*−/−*^ mice exhibit a significantly shortened OFT (Fig 7E), most likely due to a reduced number of incorporated myocardial cells derived from a diminished progenitor pool in the anterior SHF. In addition to a failure of OFT elongation, a reduced amount of Islet1 positive cells in the intercalated cushions of mutant mice (Fig. 5) cause hypoplastic endocardial cushions (Fig. 2), further impairing OFT septation. As intercalated cushions are formed from SHF-derived epithelial progenitor cells (38) that express LRP2 (Fig. 3B), this receptor is likely important for efficient migration of Islet1 positive epithelial SHF progenitor cells into the intercalated cushions.

### LRP2 deficiency impairs SHH-dependent expansion of the DPW progenitor zone

To facilitate OFT elongation, SHF progenitor cells exhibit an increased proliferative capacity and a delayed propensity to differentiate into myocardium (39). Several signaling pathways have been implicated in balancing proliferative versus differentiation fate decisions in the SHF (24, 40) -when defective, they cause CAT. These pathways include canonical and non-canonical Wnt signaling, as well as signals downstream of FGF, BMP, and SHH (27, 41–47). Our studies failed to provide evidence for defects in Wnt and BMP signaling as primary cause of CAT formation in the LRP2-deficient heart. Rather, our findings from the *Gli1_LacZ* (Fig. 6) and the *Gli1_CreER*^*T2*^ (Fig. 7A-C, 8A, B) reporter strains strongly argue for a defect in SHH signaling in the progenitor niche of the DPW as the underlying pathological mechanism. Our model is consistent with the established role of LRP2 in promoting SHH signal reception in other embryonic tissues, such as the neuroepithelium (10).

The manifold cardiac phenotypes observed upon interception with the SHH pathway uncovered multiple roles for this morphogen pathway in heart morphogenesis (48). SHH is necessary for arterial pole formation as loss of SHH signaling in *Shh*^*−/−*^ mice or in chick embryos treated with the SHH antagonist cyclopamine results in CAT formation (49, 50). Conditional disruption of the SHH pathway mediator *Smoothened* in Islet1 positive progenitor cells or a *Shh* conditional mutant using the *Nkx2.5*^*cre*^ deleter strain substantiated a role for SHH in control of OFT elongation and identified SHF progenitors as cell population receptive to SHH signals. Our findings now identify LRP2 as an essential component of the SHH signaling machinery during OFT formation and implicate defects in SHH signaling in the DPW in CAT formation in receptor mutant mice. In addition to its role in arterial pole formation, SHH has also been reported to regulate posterior heart formation. One class of phenotypes are atrial and atrioventricular septal defects identified in patients and mice genetically deficient for hedgehog signaling components (51–54). These defects arise due to the importance of SHH signaling in specifying a subset of atrial progenitor cells in the posterior SHF. This subset of cardiac progenitors forms the atrial septum as well as the dorsal mesenchymal protrusion responsible for formation of the atrioventricular septal complex.

In addition to CAT formation, *Lrp2*^*−/−*^ embryos show ventricular septal as well as aortic arch defects as reported by others. Such malformations also occur in *Shh*^*−/−*^ mice and in animals defective in the ciliary signaling compartment (14, 50, 55). While it is conceivable that ventricular septal defects in *Lrp2* mutants documented before (16) may also be caused by SHH signaling defects, our study focused on the consequences of impaired SHH signaling in the anterior SHF region for CAT formation. Using the *Gli1-CreER*^*T2*^ reporter line, we show an overall reduction in the number of SHH-responsive Islet1 positive progenitor cells in the *Lrp2*^*−/−*^ distal OFT tissue (Fig. 7A, C). Also, while these cells were mainly restricted to the DPW and the Tz in controls (Fig. 8A), they showed a more dispersed localization in the OFT tissue in mutants, overlapping with MF20 positive cardiac cells in Tz and OFT myocardium. In line with an impaired demarcation of DPW, Tz, and OFT myocardium in mutants, the expression domain for cardiac troponin I reached into the Tz, while E-cadherin expression, restricted to the Tz in controls, aberrantly extended more posteriorly into the mutant DPW (Fig. 8C). Finally, the pattern of laminin, indicative of a more mature development of the basal lamina was restricted to proximal regions of the Tz and the OFT myocardium in controls but extended inside the distal Tz and anterior DPW in *Lrp2*^*−/−*^ heart tissue. Collectively, these findings suggest a role for LRP2 in establishing a SHH-dependent niche for cardiac progenitor cells in the DPW and in controlling their differentiation along their path to the OFT.

### LRP2 likely acts as agonist for SHH signaling during OFT formation

In the LRP2 mutant OFT, both the number of Islet1 positive cells receptive to SHH signals (double positive for Islet1 and YFP) and the overall expression domain of Islet1 positive progenitor cells are reduced (Fig. 5). Thus, we cannot distinguish with certainty whether (a) LRP2 deficiency directly impacts SHH signaling in progenitor cells, thereby decreasing their numbers, or whether (b) LRP2 activity is required for other aspects of progenitor cell maintenance and loss of SHH activity in this niche as a secondary consequence of progenitor cell loss. However, based on evidence provided in this study and in published work, an instructive role for LRP2 in facilitation SHH signaling in SHF progenitors seems most likely.

The SHH signaling pathway is well characterized at the molecular level. In addition to the primary SHH receptor PTCH1, several auxiliary SHH binding proteins have been identified that are necessary to activate or inhibit the pathway in a context dependent manner. This modulatory activity of SHH binding proteins has mainly been documented in the embryonic neuroepithelium (56, 57), with LRP2 being one of these essential SHH co-receptors. Dependent on the biological context, LRP2 acts as pathway activator (by facilitating cell surface binding of the morphogen) (10) or as pathway inhibitor (by directing SHH to lysosomal degradation) (11). Data presented herein argue that LRP2 acts as agonist to promote responsiveness of SHF progenitor cells to SHH signals during OFT formation. This hypothesis is supported by a recent study documenting a crucial role for SHH in activating progenitor gene expression and in inhibiting premature differentiation, thereby maintaining the progenitor cell population in the SHF (58). We also show that LRP2 ensures the correct migration pattern of SHH-responsive SHF progenitors during OFT elongation. These results are in line with findings that SHH-responsive progenitors from the anterior SHF migrate into the pulmonary vessel (51). Finally, a recent study on Gata4-regulated SHH signaling in cardiac progenitor cells documented the importance of SHH signals for SHF cell migration in mice (59). In this study, the migratory defect of *Gata4* haploinsufficient SHF progenitor cells was rescued by over-activation of the Hedgehog pathway using the constitutively activated Smo mutant *SmoM2*, reconstituting cell migration from the SHF into the OFT and ensuring proper OFT formation.

In conclusion, we document a crucial role for LRP2 in maintenance of a pool of SHH-dependent progenitors in the SHF and in ensuring their differentiative fate as they migrate into the OFT tissue. Our findings have identified a novel component of the SHH signaling machinery essential for heart development and uncovered the molecular cause of conotruncal malformations in humans and mouse models lacking this receptor.

## MATERIALS AND METHODS

### Mouse models

Generation of mice with targeted *Lrp2* gene disruption (2) or ENU-induced gene disruption (60) has been described before. The *Lrp2* gene defect was analyzed in receptor-deficient and somite-matched littermates either wild-type (*Lrp2*^+/+^) or heterozygous for the mutant *Lrp2* allele (*Lrp2*^*+/−*^). Since heterozygous animals demonstrate no LRP2 haploinsufficiency phenotype, *Lrp2*^*+/+*^ and *Lrp2*^*+/−*^ embryos were combined and referred to as control group in this study. Where indicated, the *Lrp2* deficient line was crossed with the *Gli1-CreER*^*T2*^ (JAX; Stock 007913), the *Gli1_LacZ* (JAX; Stock 008211), or the *Tcf/Lef_LacZ* (Mohamed et al., 2004) reporter strains. Also, where applicable, the *Lrp2* deficient line was crossed with the *Wnt1-Cre* line (obtained from C. Birchmeier, Max-Delbrueck-Center, Berlin) and subsequently with the R26R (JAX; Stock 003474) reporter line to generate the *Wnt1-Cre_LacZ* reporter strain. All experiments involving animals were performed according to institutional guidelines following approval by local authorities (X9007/17).

### Immunohistology

For hemotoxilin and eosin (H+E) staining, embryos were fixed overnight in 4% paraformaldehyde in PBS at 4°C, followed by routine paraffin embedding and sectioning at 10 μm thickness. For lacZ or immunofluorescence stainings, embryos were fixed for 1 - 4 hrs in 4% paraformaldehyde in PBS at 4°C. Fixed embryos were infiltrated with 30% sucrose in PBS overnight at 4°C, followed by 2 hrs incubation in 50% cryoprotectant Tissue-Tek® OCT in 30% sucrose, followed by 2 hrs incubation in 75% Tissue-Tek® OCT in 30% sucrose. Finally, the embryos were embedded in Tissue-Tek® OCT and subjected to 10 μm cryo-sectioning. For immunodetection of E-cadherin, PKC-ζ, cardiac troponin I, or laminin, embryos were dehydrated and embedded in paraffin and sectioned at 10 μm.

Immunohistochemical analysis was carried out by incubation of tissue sections with primary antibodies at the following dilutions: guinea pig anti-LRP2 (1:2000; made in-house by immunizing guinea pigs with full-length LRP2 isolated from rabbit kidney), mouse anti-AP2α (1:50; DSHB; Santa Cruz), mouse anti-Islet1 (1:50; DSHB), goat anti-Nkx2.5 (1:100; Santa Cruz), rabbit anti-GFP (1:400; Abcam), mouse anti-MF20 (1:50; DSHB), rat anti-BrdU (1:100; AbD Serotec), mouse anti-E-cadherin (1:200; BD Transduction Laboratories), goat anti-cardiac troponin I (1:100, Abcam), rabbit anti-laminin (1:200, Sigma), rabbit anti-PKC-ζ (1:200, Sigma), rabbit anti-Arl13b (1:200, Proteintech). Primary antibodies were visualized using secondary antisera conjugated with Alexa Fluor 488, 555, or 647 (1:500; Invitrogen and Jackson Immuno Research), or with Biotin-SP (1:100; Jackson Immuno Research) followed by fluorescent conjugates of streptavidin (1:500; Invitrogen). Image acquisitions were carried out using a Leica DMI 6000B inverted microscope or Leica SPE or Leica SP5 confocal microscopes.

### *In situ* hybridization

*In situ* hybridization on paraffin sections was described earlier (61). Plasmids for digoxigenin (DIG)-labeled riboprobes were kindly provided by L. Robertson (University of Oxford, Oxford; *Bmp4*), E. Sock (Institute for Biochemistry, Erlangen; *Sox9*), W. Birchmeier (Max-Delbrueck-Center, Berlin; *Islet1, Tbx1*), C. Birchmeier (Max-Delbrueck-Center, Berlin; *Sox10*), R. Kelly (Aix Marseille Université, Marseille; *Sema3c*), and S. Meilhac (Imagine – Institut Pasteur, Paris; *Wnt11*).

### Tamoxifen injection

Tamoxifen was prepared at 20 mg/ml in peanut butter oil. It was injected at a final dose of 2 mg into pregnant females at E7.5. Embryos were collected at E10.5 and fixed for 4 hrs, followed by cryo-sectioning and standard immunohistology using rabbit anti-GFP (1:400; Abcam) and mouse anti-Islet1 (1:50; DSHB) antibodies or guinea pig anti-LRP2 (1:2000; made in-house) and mouse anti-MF20 (1:50; DSHB) antibodies. The numbers of YFP positive and YFP/Islet1 double positive cells in the distal OFT vessel were counted on 5-9 coronal sections through the distal OFT vessel per embryo for a total of 4 control and 4 *Lrp2*^*−/−*^ embryos. The number of YFP/MF20 double positive cells in the OFT was counted on 5-8 sagittal sections per embryo for a total of 5 control and 5 *Lrp2*^*−/−*^ embryos.

### Measurement of OFT length

The length of the OFT was measured on 5 sagittal sections per embryo for a total of 6 control and 7 *Lrp2*^*−/−*^ E10.5 embryos using ImageJ.

### Statistical Analysis

Statistical significance was tested using the GraphPad Prism 7 software. Results comparing two groups were analyzed by Student’s *t* test (Figure 7 B, C, E; Figure 8 B). *P* values are indicated in the respective figure legends.

## Acknowledgement

We are grateful to Robert Kelly (Aix-Marseille Université, France) for helpful discussions and for critical reading of the manuscript and to Magali Theveniau-Ruissy (Aix-Marseille Université) for expert advice. Maria Kamprath, Melanie Großmann, Kristin Kampf, and Maria Kahlow provided technical assistance.

