## Supplementary Info for "LRP2 controls sonic hedgehog-dependent differentiation of cardiac progenitor cells during outflow tract formation"

### SUPPLEMENTARY FIGURE LEGENDS

**Supplementary Figure 1. LRP2-deficiency causes CAT or DORV formation on different genetic backgrounds**

Whole mounts (upper line) and coronal sections (lower line) of *Lrp2*$^{-/-}$ and control mouse hearts at E18.5 are shown. In control embryos aorta (Ao) and pulmonary trunk (Pa) are formed, while in *Lrp2*$^{-/-}$ embryos on a C57BL/6N genetic background a CAT is seen. *Lrp2*$^{-/267}$ mutant hearts on a mixed C57BL/6N/FVBN genetic background either form normal Ao and Pa or exhibit a milder form of OFT defects, known as a double outlet right ventricle (DORV). Scale bar: 750 µm.

**Supplementary Figure 2. Detection of SHF markers in wild-type and *Lrp2*$^{-/-}$ embryos**

(**A**) *In situ* hybridization for *Semaphorin 3c* (*Sema3c)* on sagittal and coronal sections of E10.5 embryos. No difference in expression patterns in the OFT are seen comparing *Lrp2*$^{-/-}$ and control embryos displayed on sagittal sections. On coronal sections, *Sema3c* expression is slightly reduced in the intercalated cushions of *Lrp2*$^{-/-}$ embryos compared with controls (arrowheads). Scale bar: 100 µm. (**B**) ISH for *T-Box transcription factor* (*Tbx1*) on sagittal sections of E10.5 embryos of the indicated genotypes. *Tbx1*, required for OFT development, displays similar expression patterns in the second heart field (asterisks) of *Lrp2*$^{-/-}$ and control embryos. Scale bar: 250 µm.

**Supplementary Figure 3. Canonical / noncanonical Wnt and BMP signaling in second heart field of control and *Lrp2*$^{-/-}$ embryos**

(**A**) Detection of lacZ activity on coronal and sagittal sections of the pharyngeal regions and OFT vessels from E10.5 *Tcf/Lef_LacZ* reporter mice, expressing (control) or lacking LRP2 (*Lrp2*$^{-/-}$). The activity of the canonical Wnt signaling pathway in the second heart field (as evidenced by lacZ activity) is comparable between *Lrp2*$^{-/-}$ and control embryos. Scale bar: 100 µm. (**B**) Upper panel *in situ* hybridization (ISH) for *Wnt11* on coronal heart sections of E10.5 embryos of the indicated genotypes. Expression of *Wnt11* in the OFT is comparable in *Lrp2*$^{-/-}$ and control embryos. Scale bar: 40 µm. Lower panel ISH on coronal E10.5 sections show similar patterns for *Bmp4* expression in the second heart field and distal OFT of control and *Lrp2*$^{-/-}$ embryos. Scale bar: 100 µm.

**Supplementary Figure 4. LRP2 deficiency does not affect development of primary cilia**

Immunohistological detection of Islet1, LRP2 and Arl13b on sagittal sections of the heart of control and *Lrp2*$^{-/-}$ embryos at E10.5. Similar numbers of primary cilia are detected in the DPW, Tz and in the OFT of *Lrp2*$^{-/-}$ and control embryos. In both genotypes Islet1 positive progenitor cells carry a primary cilium (detailed view in upper right corner of every image). Scale bar: 25 µm

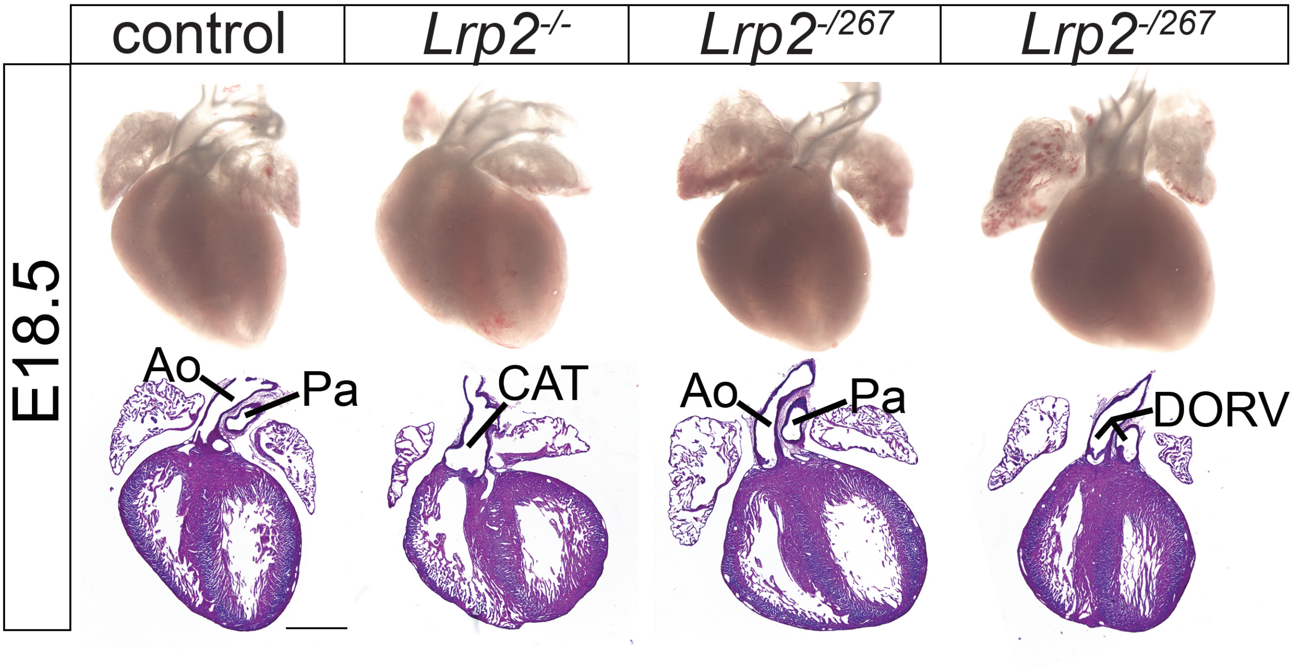


Christ and Willnow, Figure S1

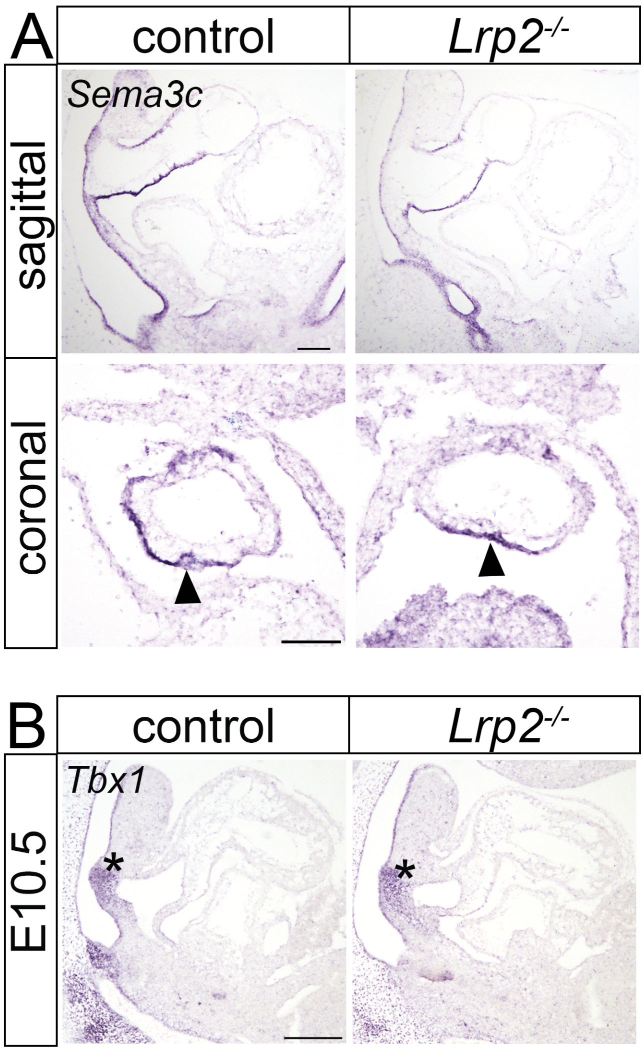


Christ and Willnow, Figure S2

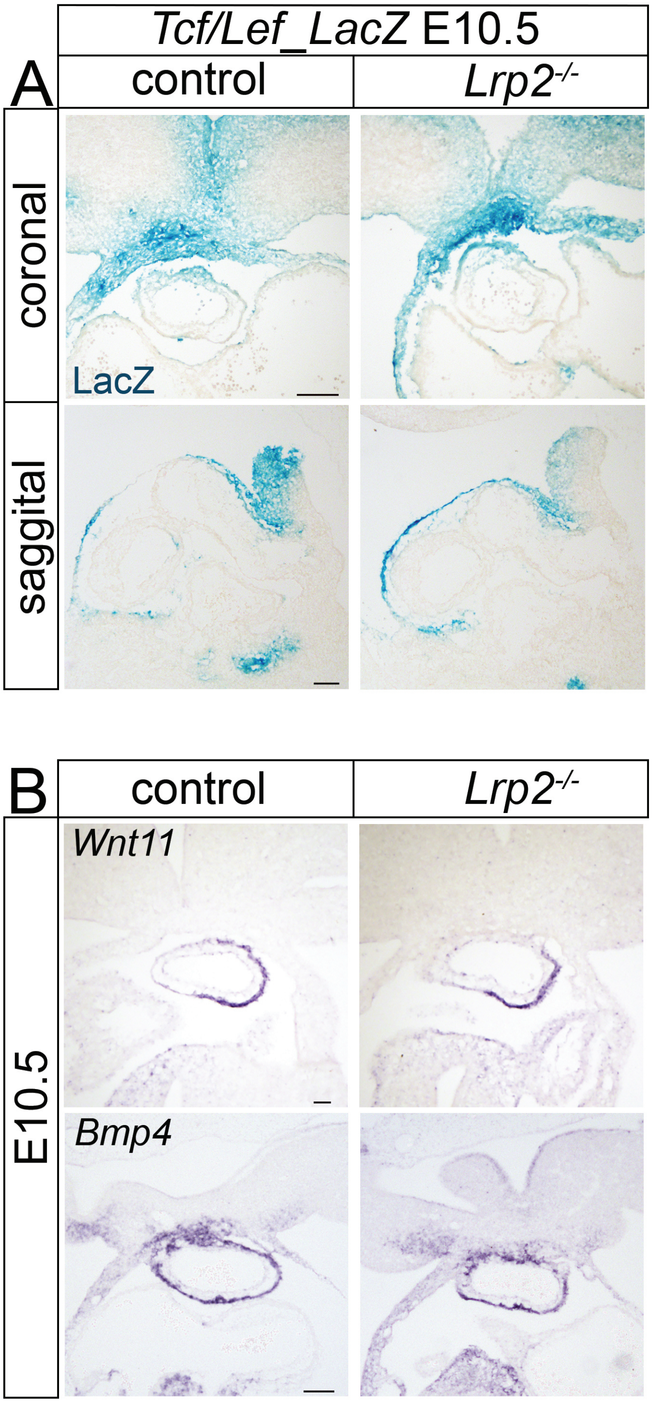


Christ and Willnow, Figure S3

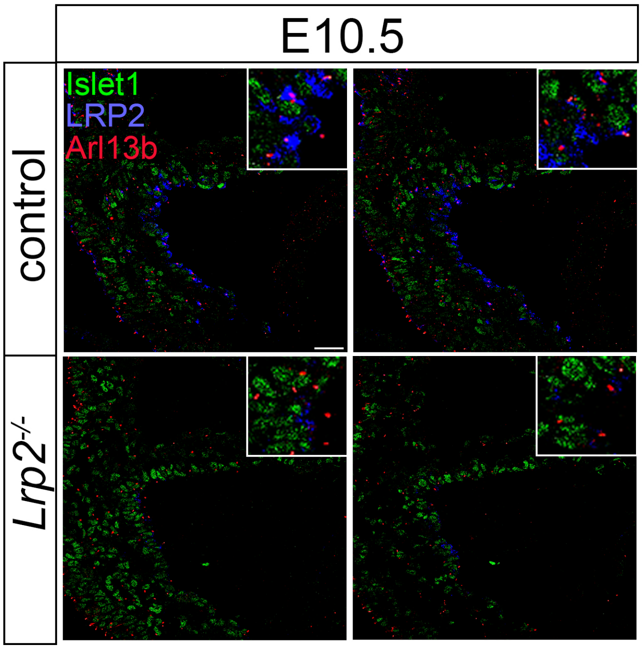


Christ and Willnow, Figure S4
